# Not so optimal: The evolution of mutual information in potassium voltage-gated channels

**DOI:** 10.1101/2020.10.26.354928

**Authors:** Alejandra Durán, Sarah Marzen

**Affiliations:** W. M. Keck Science Department, Pitzer, Scripps, and Claremont McKenna Colleges, Claremont, CA, USA

## Abstract

Potassium voltage-gated (Kv) channels need to detect and respond to rapidly changing ionic concentrations in their environment. With an essential role in regulating electric signaling, they would be expected to be optimal sensors that evolved to predict the ionic concentrations. To explore these assumptions, we use statistical mechanics in conjunction with information theory to model how animal Kv channels respond to changes in potassium concentrations in their environment. By estimating mutual information in representative Kv channel types across a variety of environments, we find two things. First, under a wide variety of environments, there is an optimal gating current that maximizes mutual information between the sensor and the environment. Second, as Kv channels evolved, they have moved towards decreasing mutual information with the environment. This either suggests that Kv channels do not need to act as sensors of their environment or that Kv channels have other functionalities that interfere with their role as sensors of their environment.

## Introduction

Accurately detecting electrical stimuli in the environment is crucial for all living organisms. It is remarkably important for communication in the nervous system, which relies on efficiently detecting and responding to electrical signals produced as neuronal ion channels open and close [1]. Such signaling is specifically dependent on the actions of voltage-gated ion channels. These are transmembrane proteins that open depending on the voltage changes across the membrane to allow an ionic current to flow [2]. Of the different types of voltage-gated channels that exist (e.g. sodium, calcium), this work focuses on potassium (Kv) voltage-gated channels, specifically those present in Metazoans (animals).

Upon voltage activation, Kv channels undergo a conformational change that allows only potassium ions to flow through [3]. They exist in all domains of life and the biological tasks they carry out are very diverse [4–6], but a roughly unified function and structure is found in Metazoan Kv channels [7]. This is due to the presence of a nervous system that sends messages in the form of repetitive current spikes called action potentials [8], which requires Kv channels to be excellent sensors. Imagine a neuron as an electrical signal passes through it: each of the voltage-gated channels needs to make the best prediction possible about the charges it feels that ultimately determine the voltage changes - in no more than two milliseconds [9]. In a nervous system that does not allow a wide range of ion concentrations, the slightest variation in the currents conveys important information about the signal, and it seems that Kv channels should accurately detect it.

Researchers have so far developed Kv channel models to assess the probability of being in an open conformation according to changes in membrane potential (voltage). Using statistical mechanics and thermodynamics is a common approach [4, 10, 11], where a variety of parameters in partition functions are introduced to generate models coherent with experimental results. The principal models of this type have been proposed by Sigworth [12] and Sigg and Bezanilla [13].

These models have been mostly used in conjunction with mutagenesis studies to determine the structure of the Kv voltage sensor domain (VSD) [14, 15] or the physiological effects of common mutations [16, 17]. However, no one has yet explored how well Kv channels sense their environment based on these models using mutual information [23]. In an environment that (it would seem) demands accurate and incredibly fast sensing, this approach is ideal to assess how informative Kv channels need to be about their environment.

In this work, we manipulate Sigworth’s model to create one that predicts how likely the channel is to be open depending ultimately on K+ concentration. Using information theory, we quantify how well Kv channels send messages about their environment. We then explore if their evolution could reflect a tendency to maximize mutual information. The following section provides a theoretical background on the biophysical model implemented, information theory, and Kv evolutionary history. In the Methods, we explain the model. In Results, we show how the gating current is a critical factor for sensing across different environments and explore the relationship between mutual information and Kv channel evolution. Finally, in the Discussion, we explore new evolutionary perspectives suggested by our results.

## Theoretical background

We first discuss models of voltage-gated ion channels, and follow that with a discussion of mutual information, our principle metric for studying the quality of a sensor.

### Modeling a two-state voltage-gated ion channel

Several biological sensors can be considered as allosterically regulated molecules, where an indirect regulator induces a conformational change [18]. Examples range from ligand-gated ion channels, to hemoglobin changing conformation upon oxygen binding. Considering these sensors as allosteric molecules allows to formulate statistical mechanical models that link changes in conformation to their external regulators, assigning statistical weights to different conformational states [19].

As for voltage-gated ion channels, a simple statistical mechanical model that describes the influence of the membrane potential (voltage difference) on the conformation of the channel was proposed by Sigworth in 1994 [12]. Assuming a model where the channel can be either open or closed, the following relation is found:

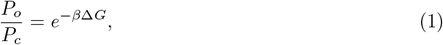

where *P*_*o*_ is the probability to be in the open conformation and *P*_*c*_ in the closed one, 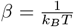 has its usual meaning where *k*_*B*_ is the Boltzmann constant and *T* the temperature in Kelvin, and

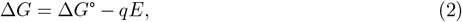

where Δ*G*° is the free energy change between the open and closed states at zero membrane potential, *q* is the gating current (in terms of elemental charges *e*^*o*^) and *E* is the membrane potential in mV. The standard Gibbs free energy Δ*G*° and *q* are unique to each type of channel. Δ*G* has been redefined by several authors [20], but the original definition is also valid. The gating current is generated when specific charges in the voltage sensor domains of the voltage-gated channel move from a lower gate to an upper gate, which is necessary for a conformational change to occur [8]. See Fig 1. This *q* value is unique for each type of channel.

**Fig 1.**
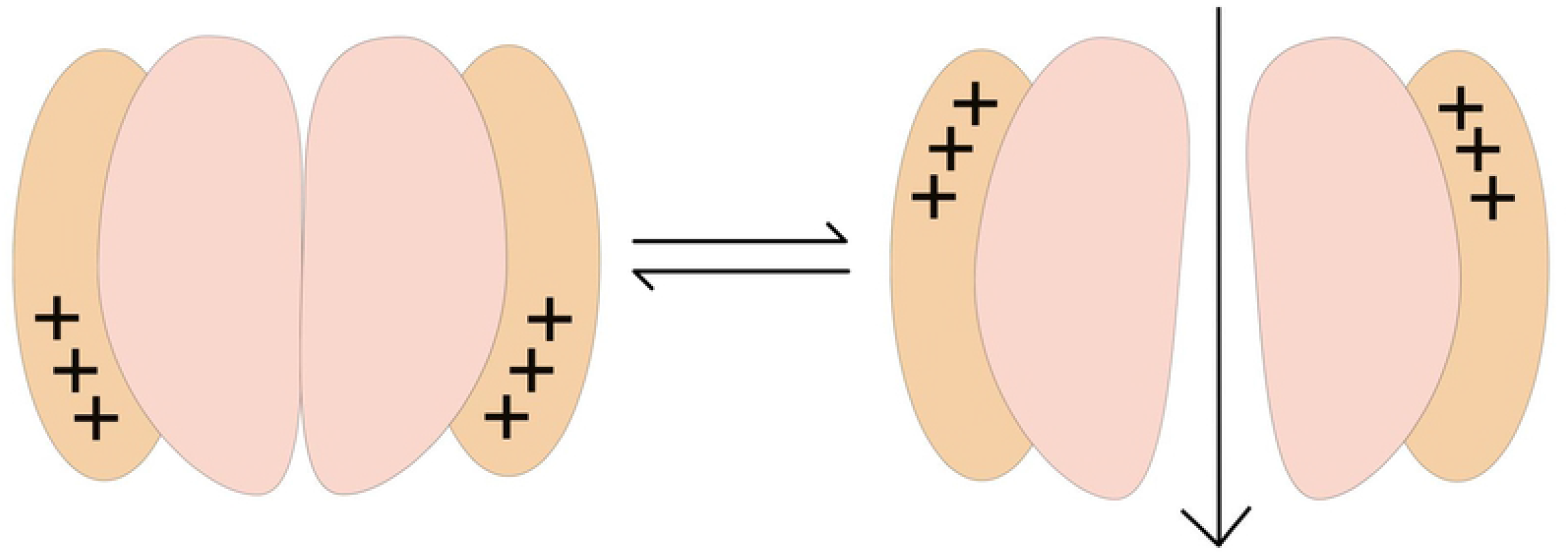
Two-state toy model of a Kv channel. The upwards movement of gating charges (in black) generates a gating current that induces the conformational change from a closed to an open state (voltage sensor domains in orange).

**Fig 2.**
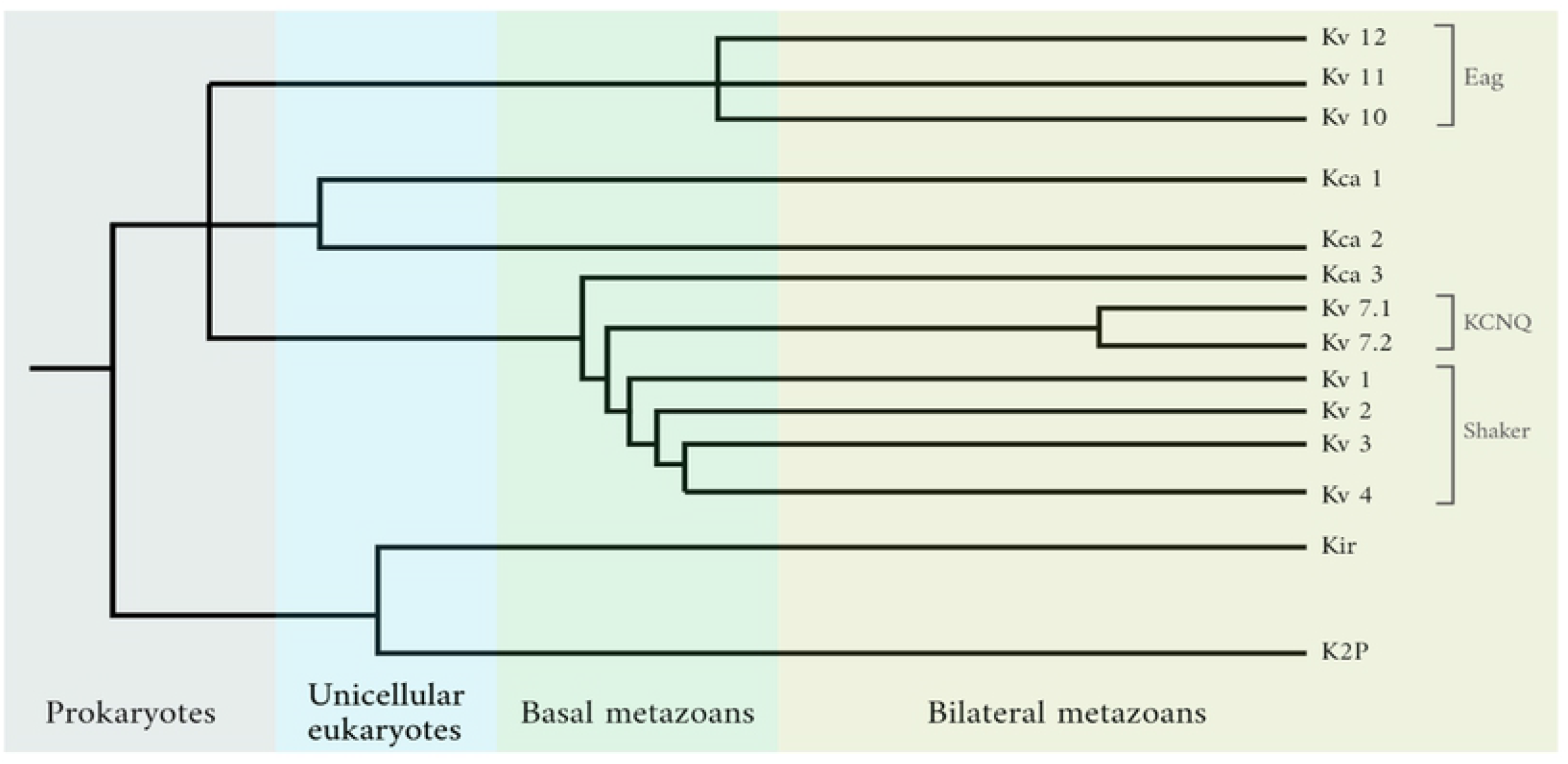
Evolution of potassium-specific ion channels during the time of appearance of 4 groups of organisms. Data taken from Li et al. [35]. Kv channel families are indicated. Unicellular eukaryotes refer specifically to the choanoflagellates. Basal metazoans include Ctenophora (comb jellies) and Cnidaria (jellyfish, sea anemones, and corals). Branch length is not proportional to time.

Knowing that the channel is either open or closed, so that *P*_*O*_ + *P*_*C*_ = 1, we find that

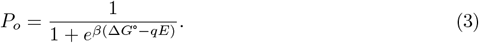

The probability of being open is affected by the gating charge and the standard free energy and is dependent on the changes in membrane potential. One can more correctly think about this as a conditional probability distribution of the channel state given the membrane potential: *P* (*open*|*E*). This notation will be used below. Later, we will relate the membrane potential to environmental concentrations of potassium, and as such, the probability distribution of channel state is conditioned on outside potassium concentration: *P* (*open* | [*K*^+^]_*o*_).

Note from equation Eq 3 that at very positive (hyperpolarized) membrane potentials, Δ*G* becomes more negative, and the channel is virtually always open. In turn, the value of *q* determines the magnitude by which changes in *E* impact Δ*G*. Generally, *q* is regarded as an indicator of the voltage sensitivity of the channel.

The model was first used by Sigworth to study the shifts in *P*_*o*_ induced by a mutation in Kv Shaker channels and has continued to be used for similar purposes [20–22]. While its simplicity makes it unsuitable for perfect fits to experimental data, it is preferred for the purposes of this work. With few and clear parameters that influence it, inferring significant contributions from each of the cases studied (section 3) is more straightforward. For the same reasons, a two-state model was chosen instead of a multi-state scheme.

### Mutual information

The model above presents a scenario where the open conformation of a voltage-gated channel is dependent on the membrane potential and thus the external potassium concentration. The open conformation immediately allows an ionic current to flow and is consequently equivalent to firing an electrical signal. In a very general sense, this can be considered as an output signal (*Y*) depending on certain input (*X*). Biology has greatly focused on how a cell or an organism gets from receiving *X* to producing *Y*, studying molecular mechanisms. However, looking at how well they process and communicate these signals in terms of the information they carry is also possible. In fact, the question about how much information from a given variable *X* can be reliably tracked to an output *Y* (or vice versa, how much can *Y* reliably tell about *X*) is answered by information theory.

Both *X* and *Y* represent random variables, meaning that their state is unknown and random until we do a particular experiment and obtain a “realization” of each. Their realizations are denoted *x* and *y*, respectively. Over the course of *N ≫* 1 experiments, we see *X* take a value *x* and *Y* take a value *y* with frequency ≈ *Np*(*x, y*). The quantity *p*(*x, y*) is known as the joint probability distribution, and it describes (equivalently, in a Bayesian sense) our belief that *X* will take value *x* and *Y* will take value *y* in any given experiment. Of particular interest are: the marginal distributions *p*(*x*) = ∑_*y*_ *p*(*x, y*) and *p*(*y*) = ∑_*x*_ *p*(*x*, |*y*), which represent the probability of seeing a particular value *x* or a particular value *y*; and the conditional probability distributions *p*(*x*|*y*) and *p*(*y*|*x*), which represent the probability of seeing a particular value *x* conditioned on the fact that we have seen a particular value *y*, or vice versa. It is of calculational importance that there is a simple relationship between joint, marginal, and conditional probability distributions: *p*(*x, y*) = *p*(*x*)*p*(*y*|*x*) = *p*(*y*)*p*(*x*|*y*). Though we have implicitly focused on the case that *x* and *y* can only be in a finite set, one can straightforwardly extend mutual information (described below) to the case when either *x* or *y* or both are real numbers.

Proposed by Claude Shannon in 1948 [23], information theory is built upon the idea of entropy in a system. This is not to be confused with the Boltzmann-Gibbs entropy used in thermodynamics. Instead, Shannon’s entropy measures the uncertainty of the state of a variable in a system (*X* or *Y* as outlined above). It reflects the average number of yes/no questions needed to correctly guess the state of a random variable [24]. It is mathematically expressed as:

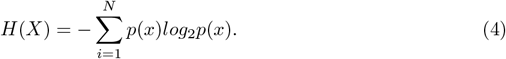

In this expression, *N* is the number of possible states of *X* (any given variable), *p*(*x*) is the probability distribution of the states of *X*, and the logarithm with base 2 allows us to get an entropy value in binary units – bits. The probability distribution can be obtained using theoretical predictive models, using statistical weights for example (as done in Sigworth’s model). As well, it can be inferred from experimental measurements [24]. For convenience, values of *X* will represent “inputs” and *Y* “outputs” throughout the rest of the paper.

To describe the amount of shared information between two systems (or two variables *X* and *Y*), Shannon introduced mutual information. It quantifies the reduction in uncertainty about *X* obtained from knowing *Y*, or vice versa. Different relationships between *X* and *Y* result in different expressions for mutual information, which have been reviewed in other works [24–26]. Here, the scenario of interest is where the state of *Y* is determined by *X*: it is conditional. The corresponding expression for its mutual information is:

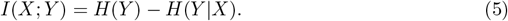

In this expression, *H*(*Y*|*X*) is a conditional entropy, which expresses the uncertainty of a certain value of *Y* occurring given *X*. It is directly related to the conditional probability distribution *p*(*y*|*x*) via *H*(*Y*| *X*) = ∑_*y*_ *p*(*y*|*x*) log_2_ *p*(*y*|*x*), which in this case, is exactly what we get using Sigworth’s model. Note the correspondence with Eq. 3. The specifications to get *I*(*X*; *Y*) are discussed in the Methods. Mutual information will quantify, in bits, how strong the (nonlinear) correlation is between *X* and *Y*. It is always non-negative, with values close to zero meaning the correlation is weak, and 0 meaning *X* and *Y* are completely independent from each other [23].

In a biological context, the value of mutual information indicates the number of possible environmental conditions (*X*) that a biological readout (*Y*) allows to distinguish. For example, if the mutual information value is 1 bit, then *Y* occurs due to one out of two possible states of X. With 2 bits, four states of X are possible; 3 bits represent eight possible states, and so on [27]. In this sense, the higher the value is for mutual information, the more states we can distinguish of *X* from knowledge of *Y*, and hence we can be less uncertain about it compared to when we did not know *Y*.

In general, information theory can be applied in two ways to biological contexts: with a source coding or a channel coding approach. The first approach assumes we can control the channel through which the information gets from *X* to *Y*, and focuses on how to compress the *X* signal into a *Y* response losing as less information as possible. The second approach assumes we can control the environment, studying how much data we can send through a noisy channel. A third approach treats the mutual information as a useful quantification of nonlinear correlations [28]. We mostly use the third approach in this work, though later address the second approach.

Mutual information has been repeatedly used to approach questions throughout a great variety of areas [26, 29, 30]. The copious number of studies using information-theoretic tools have led some authors to judge the efforts as too optimistic about the power of information theory [31]. However, in biological contexts, it can be a good indicator of how well a biological sensor tracks its input to its output. As well, careful uses of the theory have resulted in valuable new insights about a possible evolutionary principle to optimize mutual information given the energetic constraints that living organisms have to face [32].

### The evolution of Kv channels

Kv channels are present across all domains of life [7], and their functions in organisms other than eukaryotes are just starting to be understood. Archaean and prokaryotic Kv channels have been mainly used for structural modeling, and there are initial studies suggesting that they play a role in electrical signaling (only for bacteria) [33]. In eukaryotes, specifically Metazoans, three major families have been identified with further categories. The following table summarizes the classifications.

**Table 1.** Three major Kv channel families.

| Major family | Subfamilies | Kv number |
| --- | --- | --- |
| Shaker superfamily | Shaker (or KCNA) | 1.x |
|  | Shab (or KCNB) | 2.x |
|  | Shaw (or KCNC) | 3.x |
|  | Shal (or KCND) | 4.x |
| KCNQ | - | 7.x |
| Eag | Eag | 10.x |
|  | Erg | 11.x |
|  | Elk | 12.x |

Data taken from González et al. [34]

As the one shown below, phylogenetic trees showing the evolution of these Kv channels have been obtained using genomic data.

The tree shows the complete evolution for all ion channels that are selective for K+, and only the ones with the “Kv” label are voltage gated. We can see that most of the diversification including the emergence of all Kv families occurred in the basal metazoans. Surprisingly, structures are highly conserved between the time they diversified and the latest organisms in evolutionary history (for instance, humans). Although gene sequences have changed, the changes in the final expression of the protein are silent or insignificant, and major structural properties remain unchanged [36].

Such numerous diversifications in the basal metazoans yet highly conserved later lead to several questions. What could have driven such a sudden differentiation of Kv channels? What selective pressure was present during the basal metazoan era but not the following ones that slowed down Kv channel evolution? Further, looking at the deeper principles in the evolution of biological sensors using concepts from information theory, and reiterating the question we explore, did Kv channels in Metazoans evolve to maximize mutual information?

## Methods

To obtain a biophysical model for Kv channels we first identify its environment (*X*) as the extracellular potassium concentration [*K*^+^]_*o*_, and its biological readout (*Y*) as either being in an open or closed conformation. Although one could initially think that the environment should be the membrane potential, ultimately differences in ion concentrations generate such potentials and Kv channels must select only for potassium ions. Conversely, the opening and closing of the channels only generates changes in potassium concentrations. Hence, we can make an educated guess that these channels want to sense K+ concentrations in their outside (changing) environment, which in turn affect the voltage sensed.

Precisely, the Goldman-Hodgkin-Katz voltage equation (or Goldman equation) relates the membrane potential (*E*) to the main ion concentrations in and out of the cell [37, 38]:

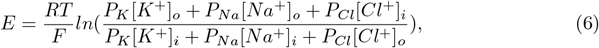

where *P* is the membrane permeability value for each ion, and the subscripts indicate if the concentration is inside (i) or outside (o) of the cell. We have implicitly assumed that the equilibrium membrane potential given the ion concentration gradients is quickly achieved.

To keep [*K*^+^]_*o*_ as the only variable term, we assume all other terms as constant depending on the respective average concentrations for each cell type. In this work, we use the concentrations for a typical neuron at rest. Then, the obtained expression for *E* can be replaced in Sigworth’s model 3 to get:

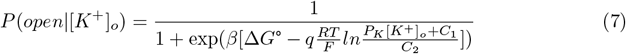

where *C*_1_, *C*_2_ are constants related to the aforementioned Goldman equation and Δ*G*° has different possible expressions. One showing high fidelity to experimental results has been proposed by Chowdhury and Chanda as the “limiting slope method” [20]. Δ*G*° values reported using this method were directly used. However, in most cases, experimental values only allowed us to make another estimation for ΔG° indicated by the same authors as:

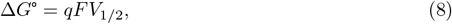

where *q* is the gating charge, *F* is Faraday’s constant, and *V*_1*/*2_ is the voltage at which half of the channels are open (half-maximal activation voltage), commonly reported in the literature.

Once we get the equation Eq 7 relating the readout (*Y*) to the identified environment (*X*), we proceed to find the expression for the mutual information. There are different ways to do so, but here we use the general expression indicated in Ref. [19]:

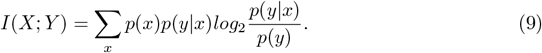

The detailed derivation can be found in the supplementary material of Statistical Mechanics of Monod–Wyman–Changeux (MWC)Models [19]. This expression is especially useful since it depends only on *p*(*x*) and *p*(*y*|*x*), which represent the known distributions of the environment and the conditional probability distribution found in Eq. 7. Recall that the marginal probability distribution *p*(*y*) is simplified because it can be expressed as:

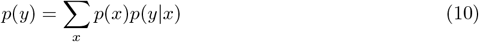

Replacing Eq 7 in Eq 9, we get the final expression for mutual information. It will quantify how good sensors Kv channels are in the context of the two-state model used.

Although it is mathematically correct for *p*(*x*) to have its domain over all real numbers, this is a condition that can never be met in real life. In this case, potassium concentrations fall within a physiological range, and exceeding it simply causes cell death. This limit varies according to cell types and tissue location. Additionally, we discretize our environment by assuming that the ion concentration can only take on a large but finite number of values within that physiological interval.

Channel capacity is defined as the maximal mutual information when *p*(*y*|*x*) is fixed and *p*(*x*) is varied– in other words, when the environment is allowed to vary, but the channel fixed. For the channel capacity calculation we use the Blahut-Arimoto algorithm [39]. Its implementation in Python along with that of all equations above is available at the GitHub repository referenced in the Appendix.

## Results

### The influence of gating current

Recalling the model obtained from Eq 7, the parameters that are expected to differentiate the responses among distinct types of Kv channels to their environment are *T*, Δ*G*° and *q*. Exploring these parameters is of interest since, should they play a significant role in the statistical mechanical model, they should also be significant for mutual information. As shown in Fig 3, changes in the gating current and Δ*G*° each impact in different ways the curves obtained. Surprisingly, changes in temperature (*T*) did not produce any noticeable changes, regardless of how greatly the value was varied, within the range 298 - 400*K*.

**Fig 3.**
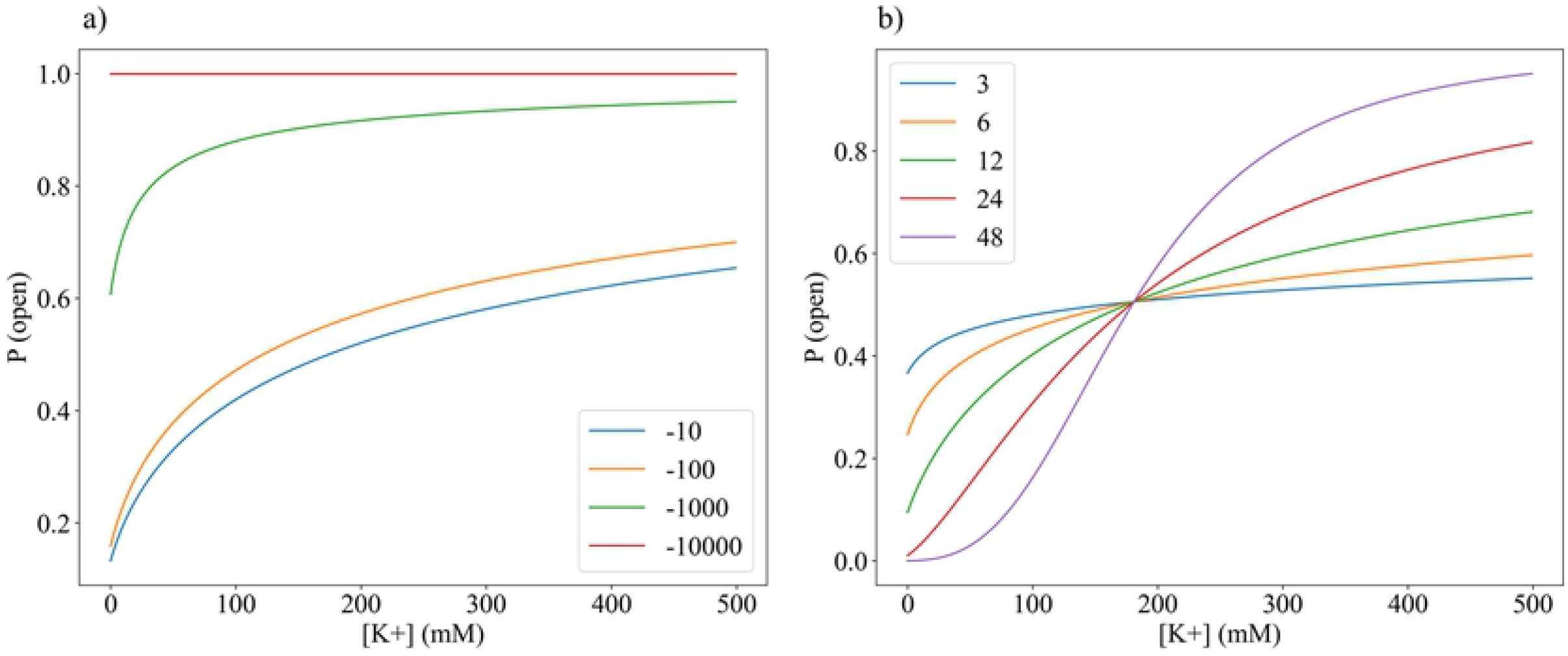
Changes in *P* (*open* | [*K*^+^]_*o*_) curves depend on standard free energy Δ*G*° with values of −10, −100, −1000, and −10000 kcal/mol (a), and increasing gating current *q* with values of 3, 6, 12, 24, and 48 *e* (b). The curves in (b) intersect at coordinate (181, 0.505).

Shifts in the shape of the curve are produced by only doubling the gating current values, while it is necessary to increase Δ*G*° by orders of magnitude. The shifts corresponding to changing the free energy are expected: the more negative Δ*G*° is, the more energetically favorable it is for the channel to be in the open conformation. However, in a biological context free energy values are very rarely below −100 kcal/mol. For comparison, the combustion of pure carbon dioxide, one of the most exergonic reactions in nature, has a free energy of −94 kcal/mol. The range of possible shapes that the *P* (*open* |[*K*^+^]_*o*_) curve could take are thus between the blue and orange curves in Fig 3, which show no outstanding variation.

However, realistic increases in the gating current cause the curve to be increasingly sigmoidal. With high gating currents, the curve starts resembling the common current-voltage (QV) curves obtained in patch-clamp experiments. Interestingly, there is a point where all curves have the same *P* (*open* |[*K*^+^]_*o*_), regardless of the gating current. This happens precisely when half of the channels are open, matching the condition that defines the median voltage of activation *V*_1*/*2_. We therefore highlight that although *V*_1*/*2_ is used to differentiate the QV curves of voltage-gated channels [8], the voltage at which half the channels are open has no dependence on gating currents, at least for Kv channels.

### Mutual information in different environments

The impact of *q* values on the *p*(*y*|*x*) functions suggest that mutual information values may also have a strong dependence on the gating current. We thus explore this dependence, but as Eq 10 shows, we must now consider the distribution of values in the environment *p*(*x*), where each x represents a value of [*K*^+^]_*o*_ in the model used. We consider four different possible distributions for it: uniform, normal, exponential, and bimodal. The goal here is not to precisely portray real-life environments, but rather to evaluate how significantly the model behavior changes just by switching to a very different distribution. Recall that under an information-theoretic source coding approach, we would expect an organism to modify itself (in this case the Kv channel) to compress data from changing environments into a signal. Then, perhaps one strategy these channels could use to do so is modifying or optimizing their gating currents, as shown in Fig 4. The mutual information values found are however significantly lower than channel capacity - the upper bound of information transmission. If mutual information was instead near channel capacity, then the channel’s sensitivity would be theoretically near-optimal, but this is not the case here (Fig 4.b).

**Fig 4.**
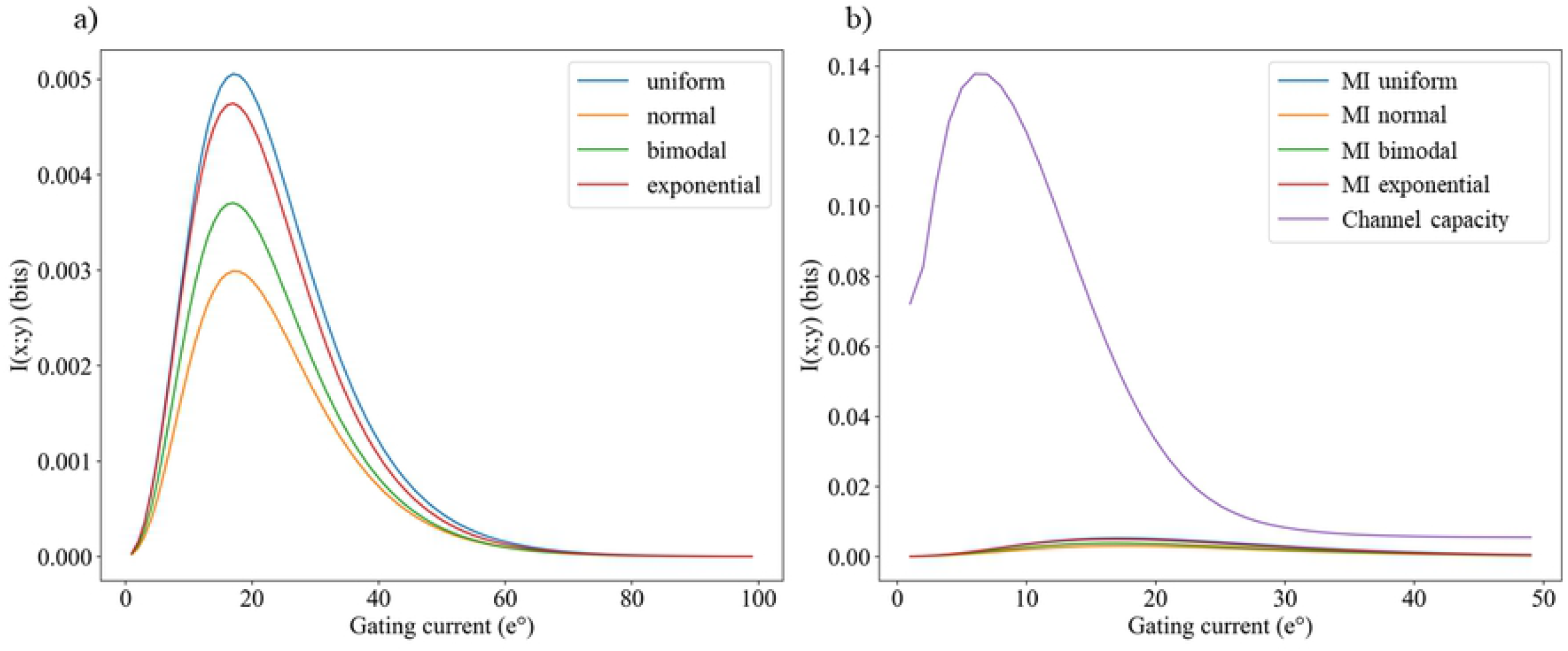
There is an optimal gating current that generates a maximum mutual information for four distinct environment distributions (a), but the values of mutual information at these optimal gating currents are far from channel capacity (b). The x (environment) ranges from 0 to 20 mM, intervals are taken each 0.5 mM, mean = 10, second mean (only for bimodal) = 3, std. deviation (for normal and bimodal) = 5. Max. mutual information values are 0.0052 (uniform), 0.0048 (exponential), 0.0037 (bimodal), and 0.0029 (normal) bits. Maximum channel capacity is 0.138.

Surprisingly, four markedly different distributions show there is a very similar optimal gating current. However, the maximum mutual information values obtained are still very low. With a uniform environment, a value of 0.005 bits tells us that if we see a Kv channel that is open, the doubts we initially have about the environment are only reduced by 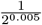, or 0.34%. It seems not to be a very suitable sensor for the task it should do, but we will return to the discussion of these values later.

It is commonly thought that the greater the gating current is, the more sensitivity the Kv channel has [40]. One could easily assert that higher sensitivity makes a better sensor. However, these results suggest that there is one value of gating current beyond which Kv channels do not improve their performance as sensors. Also, while the curves seen in Fig 4 are apparently highly conserved, no Kv channel has the gating current that would supposedly always maximize mutual information.

We also highlight that the channel capacity is remarkably low compared to those obtained in other biological systems [24, 41]. It also shows a dependence on gating current (Fig 5) with a nonzero optimal value only after certain biophysical limit of the environment.

**Fig 5.**
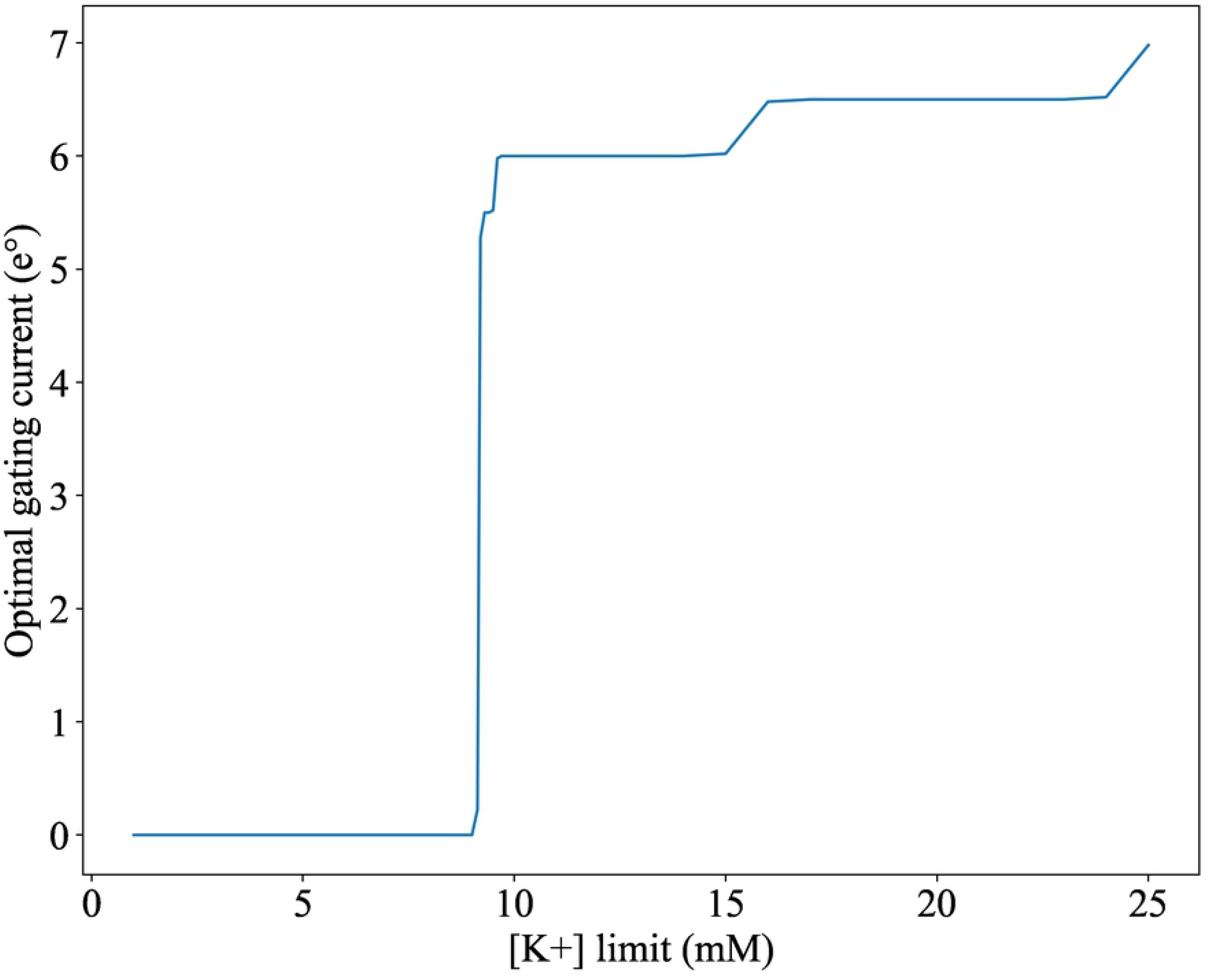
Channel capacity has optimal values depending on the [K+] limit. After the limit exceeds 9.1*mM* channel capacity has a non-zero maximum value when the gating current is initially 5.5 e°. The limit is realistically 5 *mM* on average and never higher than 10 *mM*.

### The evolution of mutual information

We now evaluate the mutual information values of Kv channels from an evolutionary approach. To have a well-rounded perspective about the different types of Kv channels that have evolved, we have chosen representative and well-studied types from the Shaker and KCNQ families (refer to Fig 2). We do not consider the Eag family because it has significant allosteric regulators other than gating currents [42], and hence does not meet the sole dependence on [*K*+] that the model used has. The considered Kv channel types with their corresponding parameters and their evolutionary order are shown below.

**Table 2.** Representative Kv channels across different families.

| Subfamily | Kv channel type | q ($e^\circ$) | $\Delta G^\circ$ (kcal/mol) | References |
| --- | --- | --- | --- | --- |
| (outgroup) | KvAP | 9.5 | -7.87 | [20, 43, 44] |
| Shaker | 1.2 | 13 | -14.63 | [44] |
| Shab | 2.1 (mammalian) | 12.5 | -10.06 | [43] |
|  | 2.1 (invertebrate) | 7.5 | -7.54 |  |
| Shal | 4.2 | 3.3 | -3.6 | [45] |
| KCNQ | 7.4 | 1.9 | -1.07 | [4] |

We identified two environments representative of specific evolutionary eras (a primitive and a recent one), to compare how mutual information evolves in different environment distributions defined also with different parameters. Although the distributions were very different, the Kv channels end up showing common behaviors.

**Table 3.** Parameters of marine and CSF environment to be used in the distribution categories.

|  | Environment parameters |  |  |
| --- | --- | --- | --- |
|  | Range [K+] (mM) | Mean | Standard deviation |
| Primitive: marine | 0 - 20 | 10 | 2.5 |
| Evolved: CSF | 3.1 - 4.0 | 3.6 | 0.3 |

The standard deviation is only used in the normal and bimodal distributions. The second mean for the bimodal is 3 in the primitive and 3.9 in the evolved environment. Data taken from Somjen [46] and Bayrami et al. [47].

We identified these environments considering the evolutionary relationships between Kv channels, which had, at most, two evolutionary hotspot moments. The first corresponds to the narrow period between the emergence of Ctenophora (comb jellies) and Cnidaria (jellyfish) during the evolutionary time of the basal metazoans Fig 2. It is known that most of the diversity of Kv channels appeared during that time, including all diversions of the baseline voltage-gated family members [35]. Although these primitive marine animals had internal nervous systems, they were very thin and did not have permeable or isolating layers. Ions were then easily diffused, making the ionic environment in their nervous system essentially identical to that of the open ocean [47]. Hence, these marine [*K*+] distributions define the first environment.

The second hotspot occurred in very late bilaterian metazoans, specifically mammals and insects. The KCNQ family greatly diverged during this time period. The environment in which they did so corresponds to the developed bilaterian nervous system with highly regulated potassium concentrations. We took the cerebrospinal fluid (CSF) as the reference to define the second environment.

Defining *p*(*x*) with these parameters allows us to use mutual information evolution to evaluate possible evolutionary scenarios. First and foremost, if Kv channels had evolved to maximize mutual information, we should expect to always see increasing curves in Fig 6. If mutual information between ion concentration and channel state had been selected for positively, the graph would show steeper slopes between the most evolved Kv channels in the CSF environment than in the marine one– mutual information would be even more maximized in the evolved environments where most recent Kv channels have been selected for. These expected trends are evidently not true. However, we highlight that a change in the steepness of the curve occurs in the normal and bimodal environments, where the most basal channel types are most greatly favored in their corresponding basal environments, although differences in mutual information are not significant in the evolved environment.

**Fig 6.**
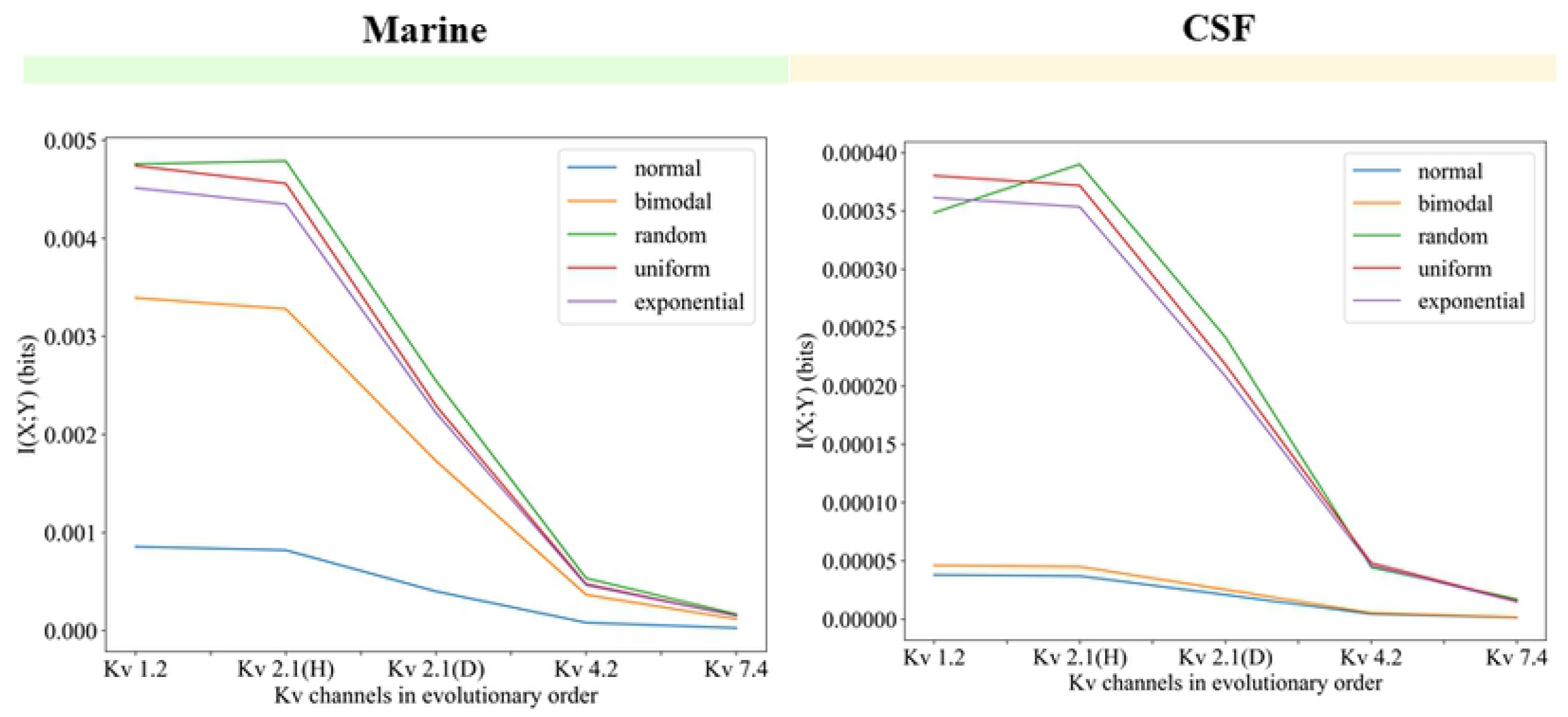
Mutual information in the evolutionary history of Kv channels. Mutual information in the evolutionary history of Kv channels always decreased in all environment distributions. The environment intervals are taken each 0.1*mM*. At the left under the green bar: real-life conditions of a primitive environment during the first evolutionary hotspot. At the right under the yellow bar: real-life conditions of an evolved environment during second hotspot. Note the color correspondence with Fig 2. Note that the random, exponential, and uniform distributions are not affected in shape or steepness when changing the parameters. Kv2.1(H) is of mammalian type (Human) and Kv2.1(D) is invertebrate (*Drosophila melanogaster*).

The decreasing trend, considering the accuracy of normally distributed concentrations [46, 48, 49], strongly suggests that maximizing mutual information was not the principle behind the evolution of Kv channels. As much as their biological task in nervous communication suggests that they should be optimal sensors tracking their environment very well in their signals, it appears not to be the case here.

Recalling Eq 5, there are two possible scenarios that may lead to decreasing mutual information values: either *H*(*Y*) has decreased or *H*(*Y*|*X*) has increased during evolutionary history. It is possible that both values changed simultaneously, and their effect decreasing mutual information shows to be correlated with the lower gating currents that emerged in Kv channels (Fig 7). The correlation is reasonable considering that gating currents are known to determine channel sensitivity and hence their ability to convey information about their environment. However, why evolution would have selected for smaller gating currents is not immediately clear. We face here a scenario where evolution appears to not care about making more efficient signals, at least under what we have considered to represent an optimal response. However, why evolution would have selected for smaller gating currents is not immediately clear. We face here a scenario where evolution appears to not care about making more efficient signals, at least under what we have considered to represent an optimal response.

**Fig 7.**
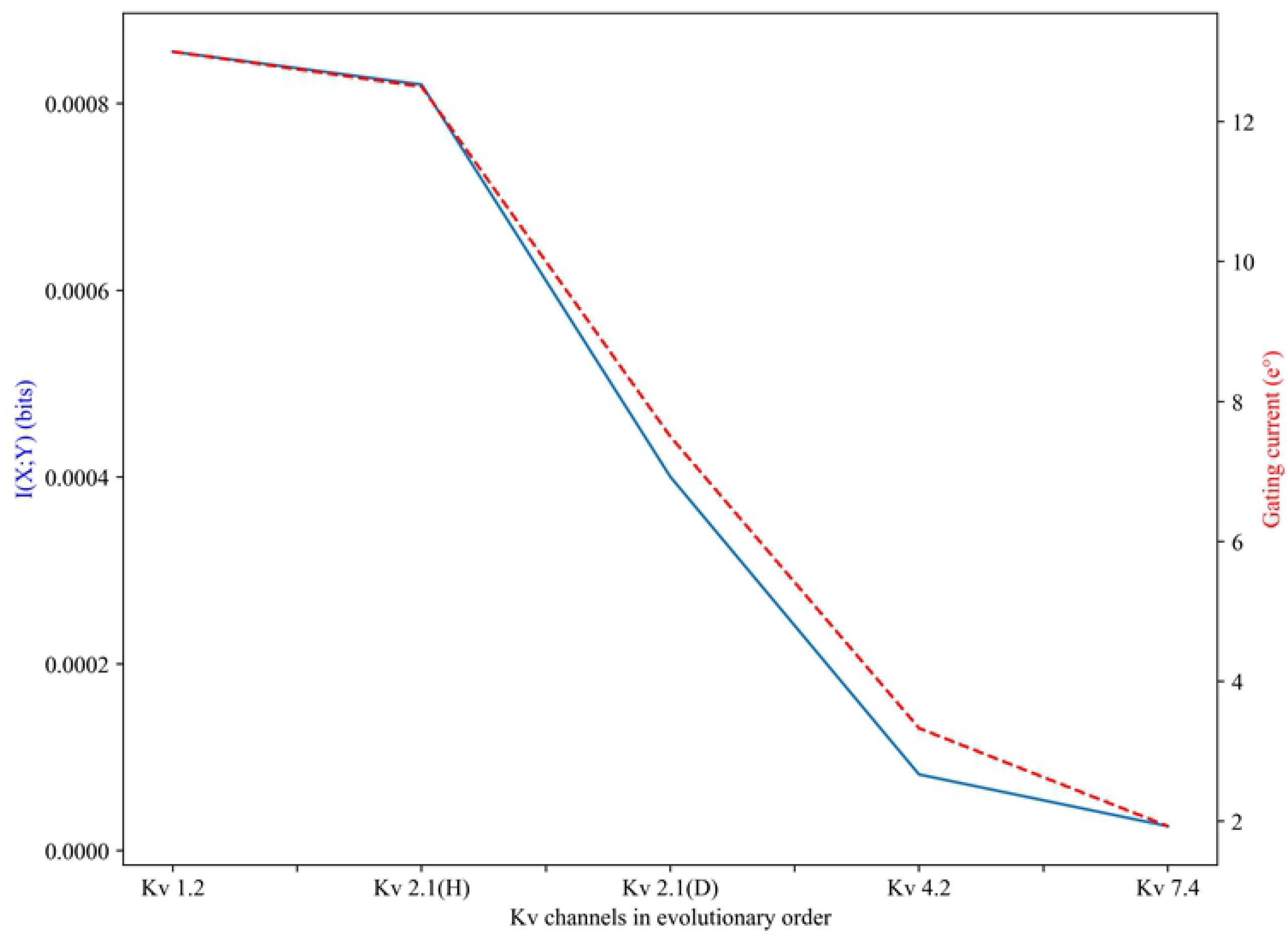
Decrease in mutual information is closely related to decrease in gating current in Kv channel evolution. The mutual information in a normally distributed environment (blue) is used as an example, but the correlation with gating current is similar in all other environments.

Looking at the more realistic normal environment, going from marine to CSF conditions also significantly decreases the maximum mutual information (*max*_*MI*_) by two orders of magnitude. Although the marine *max*_*MI*_ is still low, it is 100 times more informative than its CSF counterpart. Evidently, for two Kv channels with the same precision (e.g. they detect each 0.1*mM* of ion concentrations), the signal that responds to detecting such units will be a more specific, or less uncertain, message about the actual environment when it goes up to 20 *mM*, than another with a maximum of 4 *mM*.

In the 20 *mM* maximum case, detecting any concentration value with a 0.1 *mM* precision means that (ideally and in the simplest case) one state is differentiated from the other 199. In the 4 *mM* maximum case, one state is differentiated from only 39 other possible ones. In general, *H*(*X*) increases when the [*K*+] limit is higher, explaining why the marine *max*_*MI*_ is higher than that of the CSF environment.

Although this supports that Kv channel performance (and arguably that of other biological sensors) is affected by their precision relative to the biophysical limit of their environment, it does not show that their evolution tends to maximize the information it can get given these constraints.

## Discussion

Evaluating how good Kv channels are at accurately making predictions about their environment using an information-theoretic approach suggests that they did not evolve to maximize mutual information. Instead, the evolutionary history shows a steady decrease in mutual information that is strongly correlated to similarly decreasing gating currents. Regardless, showing what did not lead to their peculiar evolution still allows us to identify certain crucial parameters for their response, and provides insight into one of those cases where biological sensors do not need to be optimal.

It is worth highlighting that an evolutionary trend that does not seem to improve the performance of Kv channels is coherent with the high conservation seen between evolved and primitive Kv channel types. It is reasonable to suggest that during the massive diversification these channels had when basal metazoans emerged, the channels developed a sufficient amount of sensitivity and ability to predict their environment, which is surprisingly far below channel capacity. Nevertheless, there would have been no further need to maximize the acquired sensitivity, and the critical structural features were highly conserved over time.

The possibility of later diversification to be mostly random products of independent evolution has already been suggested by Anderson and Greenberg [50]. Although different channel types have different responses to macroscopic currents, they show that these variations have a negligible effect on the current that crosses when the channels open during such short action potentials. Within this context, optimizing K+ channel sensing has no real biological repercussion and then cannot be a selective pressure for evolution – a notion supported by our results.

If mutual information changes reflect random evolution, then the gating current – which we show to be highly correlated with mutual information – should also have certain randomness associated with it. However, even minimum changes in gating currents are the reason behind several nervous signaling dysfunctions [40, 42, 51].

Likewise, there is a clear trend across environments that show an optimal gating current for maximum mutual information.

Considering the ordered character of these last two facts, we suggest a possible evolutionary scenario: random diversions in protein domains (and hence gating currents) first generated functional Kv channel types, which then did not need to evolve significantly, but ended up incorporated billions of years later into such a regulated microscopic nervous “system” that random alterations to Kv channel features now have a negative impact. By this “system”, we refer to the neuron membrane complex of protein transporters and pumps actively interacting with ions, regulators, and other cells nearby. If Kv channels did not notably evolve, they generated very similar currents that ultimately regulated the ionic environment for a long time. Perhaps, Kv channels then may have represented a selective pressure to many other components in this system, which adapted to them.

This could not only explain why random variations are so impactful now but not when Kv channels emerged, but also suggest the possibility of any of the components of the system preventing the gating current to increase up to its optimal value. Possibly in a trade-off relationship, getting to the optimal gating current destabilizes a feedback mechanism or any other transduction pathway not clear yet.

As a whole, we have to acknowledge that with the model we have used, just as in any other case that models a biological situation approaching it from information theory, we can never be sure that we have correctly identified what the organism (or in this case the Kv channel) wants to sense. Perhaps Kv channels are just not well thought-of as sensors. There may be further dependencies for potassium concentrations, and even significant feedback loops or cooperativeness in the Kv channel subunits. As well, the two-state model itself may behave differently than a multi-state one. These are left as potential considerations for continuation of this work.

## Conclusion

Biological sensors do not always need to maximize mutual information, and we have shown that so is the case for potassium voltage-gated channels. Their evolution seems to have been driven by numerous random diversifications when basal animals appeared, which left no need to further optimize their performance as sensors. Still, the gating current is most likely a determinant feature for how well Kv channels sense. We find conserved tendencies for a possibly optimal gating current which still no Kv channel has, leading to possible evolutionary scenarios that may have caused this.

These tendencies are even kept when the environment distribution changes, suggesting that Kv channels could know how to keep the level of efficacy as sensors that is just good enough for them. While they cannot control their environment, they respond accordingly to how much surprise they can find in it, and opt for a same range of parameters to perform well given these constraints.

## Supporting information

**S1 Appendix**. GitHub repository URL with the code used:

https://github.com/aduranu/Kv_channel_modeling.git

## Acknowledgments

S. Marzen acknowledges support from Air Force Office of Scientific Research under award number FA9550-19-1-0411. A. Duran and S. Marzen acknowledge support from the Pioneer Research Program.

